# A Deep Learning Semiparametric Regression for Adjusting Complex Confounding Structures

**DOI:** 10.1101/2020.06.08.140418

**Authors:** Xinlei Mi, Patrick Tighe, Fei Zou, Baiming Zou

## Abstract

Deep Treatment Learning (deepTL), a robust yet efficient deep learning-based semiparametric regression approach, is proposed to adjust the complex confounding structures in comparative effectiveness analysis of observational data, e.g. electronic health record (EHR) data, in which complex confounding structures are often embedded. Specifically, we develop a deep learning neural network with a score-based ensembling scheme for flexible function approximation. An improved semiparametric procedure is further developed to enhance the performance of the proposed method under finite sample settings. Comprehensive numerical studies have demonstrated the superior performance of the proposed methods as compared with existing methods, with a remarkably reduced bias and mean squared error in parameter estimates. The proposed research is motivated by a post-surgery pain study, which is also used to illustrate the practical application of deepTL. Finally, an R package, “deepTL”, is developed to implement the proposed method.

## 1. Introduction

The amount of electronic health record (EHR) data has expanded rapidly (Shah and Tenenbaum, 2012; Murdoch and Detsky, 2013; Psaty and Larson, 2013), and is inevitably used in various data-driven analyses in health care (Chen et al., 2013). EHR data typically contain a large number of samples and often reflect daily clinical practice to offer valuable information on intervention efficacy under practical settings. Though comparative effectiveness analysis could be easily performed with randomized controlled trials (RCTs) (Britton et al., 1997; MacLehose et al., 2000; Benson and Hartz, 2000), in practice, RCTs cannot always be conducted for a variety of reasons (McCulloch et al., 2002; Curry, Reeves and Stringer, 2003). EHR data, on the other hand, are often readily available with rich information, and serve as cost-effective alternatives to RCTs (Miriovsky, Shulman and Abernethy, 2012). However, the dependence among the treatment assignment, response, and baseline characteristics can result in complicated confounding issues which can lead to biased estimation of intervention efficacy and misleading conclusions if they are not handled properly. In this paper, we aim to perform valid comparative effectiveness analysis for EHR data with complex confounding structures.

In comparative effectiveness analysis, a commonly used method to adjust for confounding factors is propensity score (PS) based methods (Rosenbaum and Rubin, 1983), including matching, covariate adjustment, stratification and inverse probability weighting (IPW) by PS. The PS methods use propensity scores to mimic RCTs such that samples with similar propensity scores have similar baseline features, and thus are frequently used to analyze EHR data (Toh, García Rodríguez and Hernán, 2011; Kazley and Ozcan, 2008). Under the strongly ignorable treatment assignment assumption, as shown by Rosenbaum and Rubin (1983), an unbiased estimate of the true treatment effect can be obtained by any of the PS-based methods. However, all PS-based methods heavily depend on the accuracy of the PS estimates, especially for PS-IPW and PS covariate adjustment (Austin, 2011).

For EHR data, confounding variables can impact the outcome and treatment allocation process in different ways with unknown functional formats, which makes PS modeling challenging. The motivating example in this paper is a post-surgery pain EHR data set (Tighe et al., 2016). One of the study objectives is to compare the effectiveness of two anesthetic procedures, nerve block versus general anesthesia, for relieving post-surgery pain intensity. Traditional methods, including a simple ANOVA analysis, a multivariate linear regression, and a PS covariate adjustment method with PS estimated by a logistic regression, all lead to a non-significant difference between the two anesthesia groups (Table 3). However, previous findings in closely related postoperative pain studies under RCT designs suggest that the two groups are significantly different (Tverskoy et al., 1990; Shir, Raja and Frank, 1994). This raises awareness of the possibility that traditional parametric methods may fail to adequately detect complicated structures describing the connections among the pain intensity, the anesthesia and the covariates.

The performance of traditional parametric statistical methods heavily depends on their assumptions, such as the linearity assumption in the least square regression or logistic regression. To deal with the potential nonlinearity and other complexity in EHR data, non-parametric methods can be applied, such as kernel-based methods, including Nadaraya-Watson kernel estimators, Gaussian process models (Williams and Barber, 1998), and kernel-based support vector machines (SVMs). However, these local kernel-based machines are sensitive to the curse of dimensionality (Bengio, Delal-leau and Roux, 2006). Though SVMs suffer less from the increase of dimensionality due to sufficient regularization and post-processing techniques for discrete outcomes (Platt et al., 1999), it is argued that they may not be reliable for binary or multinomial outcomes (Tipping, 2001). In the recent literature, the predictive modeling techniques for comparative effectiveness analysis have expanded, and now include Lasso, gradient boosting machine, random forest and neural networks (Chernozhukov et al., 2016; Nie and Wager, 2017; Chernozhukov et al., 2018).

Apart from the PS framework, another commonly used strategy in comparative effectiveness analysis is to employ a semiparametric framework, as given below,

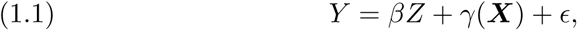

where (*Y, Z*, ***X***) is the vector of the outcome, binary treatment assignment status, and observed covariates; *β* is the treatment effect, *γ* is an unknown continuous function of ***X***, and *ϵ* ∼ N(0, *σ*^2^). This semiparametric model has been widely investigated in the statistical literature (Engle et al., 1986; Robinson, 1988; Stock, 1991). Robinson (1988) proposed an innovative strategy for obtaining an estimate of *β* with an optimal root-N-convergence rate. Instead of modeling Equation (1.1) directly, Robinson (1988) proposed the following semiparametric model,

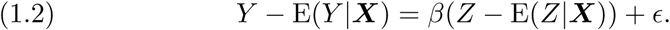

With E(*Y* |***X***) and E(*Z*|***X***) pre-estimated from the Nadaraya-Watson kernel machine approach, a root-N-consistent estimate for *β* can be obtained via a simple linear regression model based on Model (1.2) (Robinson, 1988).

Despite its root-N-consistency, Robinson’s estimator has several limitations when applied to real-world EHR data. The performance of this semi-parametric modeling strategy heavily depends on the accuracy of the estimation of E(*Y* |***X***) and E(*Z*|***X***). One obvious drawback is that the prediction accuracy of the Nadaraya-Watson kernel approach drops dramatically with the increase of the number of covariates due to the curse of dimensionality (Bengio, Delalleau and Roux, 2006). In this paper, we employ the deep neural network (DNN), a fully-connected and feedforward neural network with multiple hidden layers, as a function approximator in the proposed framework of Robinson (1988) for comparative effectiveness analysis. The neural network, a universally consistent function approximator, can approximate continuous functions on compact sets under certain assumptions (Cybenko, 1989; Faragó and Lugosi, 1993). The strong universal consistency of the neural network offers a great potential to model complicated data, compared to traditional methods such as logistic regression (Tu, 1996). To improve the accuracy and address the potential overfitting issue of the DNN, we further develop a score-based ensembling scheme via bootstrap aggregating (bagging) (Breiman, 1996).

Moreover, in modeling E(*Y* |***X***) *≡ ξ*(***X***), the residual *Y* − E(*Y* |***X***) = *β*(*Z* − E(*Z*|***X***)) +, no longer follows a unimodal distribution, especially when *β* deviates far from 0, which can potentially lead to inefficient estimation of *ξ*(***X***). To minimize the impact from these limitations and offer an accurate estimate of the treatment effect, we propose a revised semiparametric procedure.

The rest of the paper is arranged as follows. A brief introduction of DNN implementation and an improved DNN ensemble model will first be presented in Section 2, followed by a detailed illustration of the modified semiparametric framework: deep treatment learning (deepTL). Extensive simulation studies and an application of the proposed method to a post-surgery pain study are presented in Section 3. Discussions and remarks conclude the paper in Section 4.

## 2. Deep Treatment Learning

### 2.1. Deep Neural Network

For a general review of DNNs, we refer the readers to LeCun, Bengio and Hinton (2015). In this paper, we only introduce minimum but necessary concepts, to facilitate the description of the proposed method.

We first introduce the general form of an *L*-hidden-layer feedforward DNN. The model contains *L* hidden layers of nodes that transform the initial input covariates ***X*** to the estimation of the output *R*, which can be a continuous or a binary response. For example, in our semiparametric regression framework, to model E(*Z*|***X***), *R* = *Z* is a binary treatment assignment, while modeling E(*Y* |***X***), *R* = *Y* is a continuous response. For each hidden layer *l* ∈{1, … *L*}, the model takes the input, denoted by ***h***^(*l*−1)^, from the previous layer and the outputs ***h***^(*l*)^ through function ***g***_*l*_(***X***). For any *l*, denote *n*_*l*_ as the number of elements in ***h***^(*l*)^, then

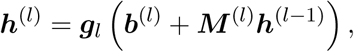

where ***b***^(*l*)^ is the bias vector with length *n*_*l*_ and ***M*** ^(*l*)^ is an *n*_*l*_ *× n*_*l*−1_ weight matrix. The function ***g***_*l*_ is regarded as applying an activation function *g*_*l*_ element-wise to the *n*_*l*_ dimensional vector ***b***^(*l*)^ + ***M*** ^(*l*)^***h***^(*l*−1)^. The activation function *g*_*l*_ is a non-linear function that transforms the output values of neurons in the previous layer into the input values of the next layer. Often a common function *g* is applied to all *g*_*l*_’s (*l* = 1, …, *L*), e.g. a rectified linear unit (ReLU) function (Hornik, Stinchcombe and White, 1989). For the first layer, i.e. *l* = 1, ***h***^(0)^ is simply the original *p* dimensional feature ***X***, and ***g***_1_ takes a *p* dimensional input and produces an *n*_1_ dimensional output. Finally, the *L*^th^ hidden layer ***h***^(*L*)^ is tied to the output *R* through

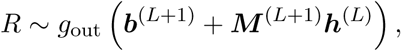

where *g*_out_ gives a scalar output.

The output function is selected depending on the outcome *R*. For a continuous *R*, we use an identity output function *g*_out_(*t*) = *t*; while for a binary *R*, a sigmoid output function *g*_out_(*t*) = 1*/*(1 + *e*^−*t*^) is used.

The final convolved output function is

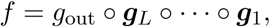

then *f* takes ***X*** as input and contains 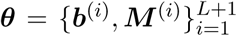 as a collection of parameters. Under this setup, ***θ*** can be estimated by minimizing the following empirical risk function,

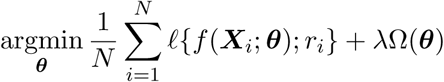

where *ℓ*(·; ·) is the loss function, Ω(***θ***) is a penalty function and *λ* is a hyperparameter that controls the degree of regularization. For a continuous *R*, we set the loss function *ℓ*(*f, r*) to 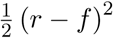. For a binary *R, ℓ*(*f, r*) is set to the Bernoulli negative log-likelihood, − {*r* log *f* + (1 − *r*) log (1 − *f*)}. Furthermore, to shrink the model size, we put an *l*_1_ regularizer on the weight matrices ***M*** ^(*l*)^ (*l* = 1, …, *L* + 1), such 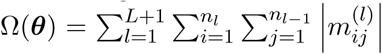 is used, where 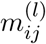 is the *ij*^th^ element in the weight matrix ***M*** ^(*l*)^.

We optimize the risk function by using the mini-batch stochastic gradient descent algorithm (Byrd et al., 2012), together with an adaptive learning rate adjustment method, i.e. adaptive moment estimation (Adam) (Kinga and Adam, 2015).

### 2.2. Bootstrap Aggregating

The total number of parameters in ***θ*** is 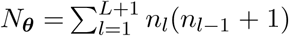, which is usually greater than the sample size *N*, leading to over-parameterization and unstable prediction. The accuracy of the prediction from a single DNN model, therefore, is expected to be unreliable when the sample size is finite. An ensemble of neural networks has been shown to outperform a single neural network (Hansen and Salamon, 1990) in such scenarios. Thus, we apply bagging, i.e. bootstrap aggregating (Breiman, 1996), to increase the robustness and accuracy of DNNs.

Specifically, we randomly sample the training set with replacement *K* times (i.e., bootstrap samples). Each time we fit a DNN model using a bootstrap sample and keep all the unused samples, namely the out-of-bag (OOB) samples, as the validation set. Let 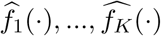 be the fitted models out of the *K* bootstrap samples. To aggregate the predictions from the *K* fitted models, a natural choice of the aggregated bagging prediction for a new observation with input ***x*** is 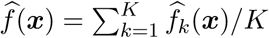.

However, due to the randomness of the initial parameters, some DNNs may not converge to a stable solution and thus perform poorly. In neural network ensembles, it is argued that “many could be better than all”, meaning that using a subset of bagged DNNs that are well fit to the data could be better than using all of them (Zhou, Wu and Tang, 2002; Mi, Zou and Zhu, 2019). Therefore, in our proposed method, we remove certain DNNs according to criteria defined below, which is consequently beneficial to the final ensemble model. For the *k*^th^ bootstrap sample, we define a performance score *v*_*k*_ as follows,

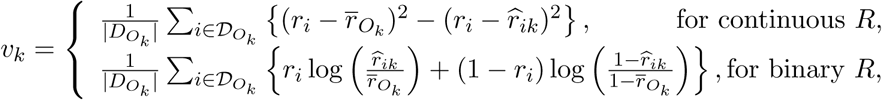

where 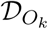 is the set of OOB samples with 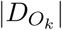 being the associated sample size, 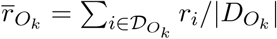 and 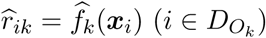. For the regression DNN, *v*_*k*_ is the mean squared error loss, while for the classification DNN, *v*_*k*_ can be interpreted as the negated binomial deviance.

To determine the optimal subset of DNNs retained in the ensemble, we first rank DNNs based on their performance scores, i.e. *v*_(1)_ ≥ … ≥ *v*_(*K*)_. The prediction for ***x***_*i*_ (*i* = 1, …, *N*) by aggregating the top *q* DNNs is then,

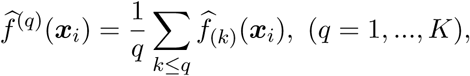

where 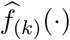 is the fitted DNN corresponding to the performance score *v*_(*k*)_. The optimal number of DNNs utilized by the ensemble, *q*_opt_, is determined by minimizing the training loss, i.e.,

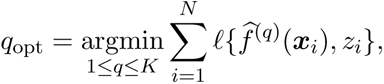

based on which we obtain the revised bagging prediction 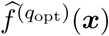 (***x***) for a new observation with input ***x***.

### 2.3. Revised Semiparametric Regression

Following (1.2), we let ξ(***X***) *≡* E(*Y* |***X***) and *e*(***X***) *≡* Pr(*Z* = 1|***X***) = E(*Z*|***X***). As *Z* in this paper is a binary treatment assignment status, *e*(***X***) is the conditional probability for a subject being assigned to the treated group, or the PS in Rosenbaum and Rubin (1983). Let (*Y*_*i*_, *Z*_*i*_, ***X***_*i*_) be the observed data of the *i*^th^ sample (*i* = 1, …, *N*), and 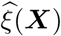 and *ê* (***X***) be the corresponding approximation functions of ξ(***X***) and *e*(***X***), respectively, estimated from the observed data. Then, we can rewrite (1.2) as:

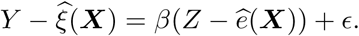

The estimate of *β* and its associated variance estimation can be obtained from the above simple linear regression model without an intercept as follows (Robinson, 1988)

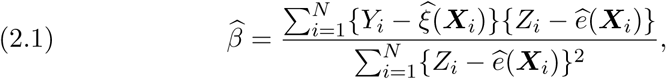

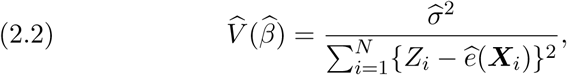

where 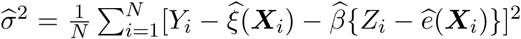. Estimate 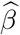 is shown to be root-N-consistent under mild conditions, i.e., 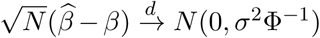and 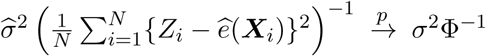, where Φ = E{*Z* − *e*(***X***)}2 (Robinson, 1988).

With an infinite number of observations from (*Y, Z*, ***X***), 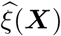 and *ê* (***X***) can consistently approximate ξ(***X***) and *e*(***X***) due to the universal consistency of DNNs (Hornik, Stinchcombe and White, 1989; Faragó and Lugosi 1993; Sonoda and Murata, 2017). However, under finite-sample situations, the model errors, 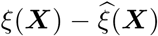 and *e*(***X***) − *ê*(***X***) exist. Note, the residual *Y* − ξ(***X***) = *β*(*Z* − *e*(***X***)) +, which is the sum of *β*(*Z* − *e*(***X***)) and. When *β* is small, the sum is dominated by, which is Gaussian, and minimizing the mean squared difference of ξ(***X***) from *Y* is equivalent to maximizing the log-likelihood of Gaussian random variables. This is expected to be efficient. However, when *β* is large, the distribution of the sum departs from a Gaussian distribution and is no longer unimodal, thus estimating ξ(***X***) by minimizing the mean squared difference can become less efficient.

Accordingly, we propose to approach Model (1.1) by replacing (1.2) with the following modified model

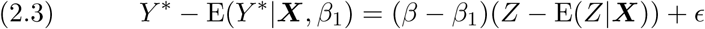

where *Y* ^*^ = *Y* − *β*_1_*Z* for a given constant *β*_1_. Instead of approximating ξ(***X***), we approximate ξ^*^(***X***, *β*_1_) = E(*Y* ^*^|***X***, *β*_1_) by our proposed score-based bagged DNNs in Section 2.2. The modified estimate of *β* and its associated variance estimation then become

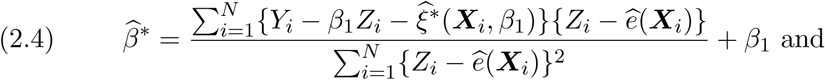

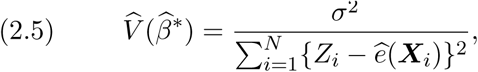

where 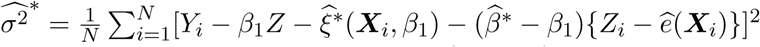.

Intuitively, if *β*_1_ is close to *β*, or when |*β* − *β*_1_| is small, the function approximation by bagged DNNs is expected to be more efficient which subsequently leads to an improved estimate of *β*. A reasonable candidate for *β*_1_ can be an estimate of *β* from any PS approach in Rosenbaum and Rubin (1983). For the simulated data and real data analysis, we set *β*_1_ to the estimate derived from the PS covariate adjustment, where the estimated PS, *ê* (***X***) from bagged DNNs, is used. The accuracy of the estimated PS, *ê*(***X***) does not depend on *β*, nor the PS covariate adjustment which only depends on *ê* (***X***) (Rosenbaum and Rubin, 1983).

In summary, deepDL first approximates *e*(***X***) by our proposed DNNs, based on which the treatment effect estimate is obtained from the PS covariate adjustment, and *β*_1_ is subsequently set to this estimate. Next, we approximate ξ^*^(***X***, *β*_1_) again by our proposed DNNs, and obtain 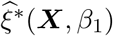. With *ê* (***X***) and 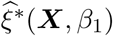 obtained, we finally get 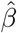 and its associated variance through (2.4) and (2.5). The modified estimator has the advantages of both the PS based method and the original semiparametric regression framework. The final algorithm is summarized as follows:

#### Algorithm 1 deepTL: A Revised Semiparametric Regression

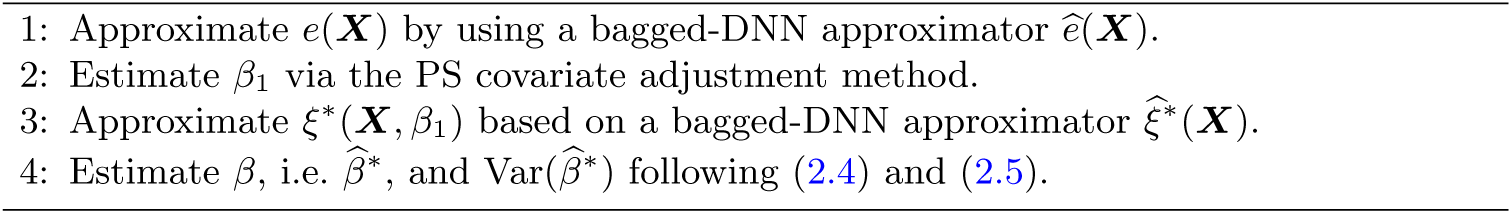

## 3. Numerical Studies

To investigate the performance of our proposed methods, we (i) conduct extensive numerical studies under different confounding structures, from simple to complex settings; and (ii) perform a real data analysis on a post-surgery pain data.

### 3.1. Simulation Studies

We conduct simulation studies in three scenarios with different confounding structures. Specifically, we adopt the following models to generate data with three confounding structures under the semiparametric framework: ***X*** ∼ MVN(**0**, *I*_*p*_), *Z*|***X*** ∼ Bin(1, *e*(***X***)), *Y* = *βZ* + *γ*(***X***) + *ϵ, ϵ* ∼ N(0, *σ*^2^) and,

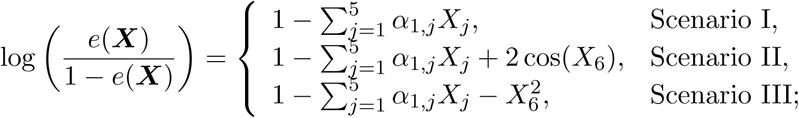

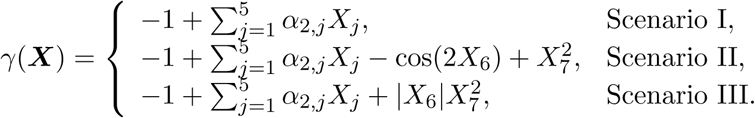

The total sample size is set to *N* = 5000, and the dimension of ***X***, *p* = 20 with the treatment effect *β* = 1 or 2, the noise size *σ*^2^ = 1. ***α***_1_ and ***α***_2_ are both 5-valued vectors with elements *α*_*k,j*_ ∼ *U* (−1, 1) i.i.d for *k* = 1, 2, *j* = 1, …, 5, which are generated at the beginning of Monte Carlo simulations and kept unchanged in the subsequent replications. All results are based on 1000 replicates. Besides the input and the output layers, all DNNs have six hidden layers, with 20, 18, 16, 14, 12 and 10 hidden nodes from the first to the last hidden layers, respectively.

Besides deepTL, we also add semiDNN, which uses the same bagged DNNs as deepTL, but directly implements Robinson’s original semiparametric procedure (Equation 2.1 and 2.2), or essentially deepTL with *β*_1_ being set to 0. In addition, we include an oracle linear regression method with underlying structures of E(*Y* |*Z*, ***X***) known, denoted as “LM-Oracle”. We also add “LM-Naive”, a multiple linear regression method with all linear terms of the observed covariates and the treatment assignment status. We further include a PS covariate adjustment method, with the PS estimated by a logistic regression, denoted as “PS-Naive”. Finally, the two cross-fitting double machine learning estimators introduced in Chernozhukov et al. (2016) are included, with all functions estimated by random forests, denoted as “DML-PLM” and “DML-DR”. DML-PLM follows Robinson (1988) and adopts the original semiparametric regression framework, while DML-DR employs a double robust framework (Robins, Rotnitzky and Zhao, 1994).

For DNNs, an *l*_1_ penalty with weight *λ* = 1*e* − 4 is applied. The mini-batch stochastic gradient descent has a batch size *N*_*B*_ = 100, and Adam is employed with a starting learning rate of 0.001. The maximum number of epochs in DNN optimization is set to 250. The bagging size *K* = 100. All random forest models consist of 2,000 trees.

To evaluate the performance of all methods, we provide the mean treatment effect estimate 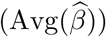, Monte Carlo standard error 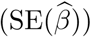, estimated standard error 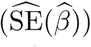, squared root of mean squared error 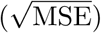 and the 95% confidence interval (95% CI) coverage.

Simulation results are summarized in Table 1. In Scenario I, because all the components are linear, LM-Naive is essentially the same as LM-Oracle. In this scenario, LM-Naive, PS-Naive, deepTL and LM-Oracle all estimate *β* unbiasedly. However, PS-Naive shows a 95% CI coverage of 98%, which is due to the inflated variance estimate of 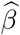 (Zou et al., 2016). In addition, we observe a small bias in DML-PLM when true *β* = 2, and thus a slightly increased MSE and a lower CI coverage than the targeted 95% coverage level. DML-DR appears to have a larger bias and Monte Carlo standard error, and the largest MSE among all the methods. Among the two bagged DNN approaches, the coverage of 95% CIs provided by deepTL is more precise than that of semiDNN. As the true *β* increases, we observe a larger bias and a remarkable decrease in 95% CI coverage in semiDNN, while the performance of deepTL remains almost unchanged, with unbiased estimate and the targeted 95% CI coverage.

**Table 1.** Simulation Results for Scenario I, II and III.

| Method | $\beta$ | $\text{Avg}(\hat{\beta})$ | $\text{SE}(\hat{\beta})$ | $\widehat{\text{SE}}(\hat{\beta})$ | $\sqrt{\text{MSE}}$ | 95% CI | $\beta$ | $\text{Avg}(\hat{\beta})$ | $\text{SE}(\hat{\beta})$ | $\widehat{\text{SE}}(\hat{\beta})$ | $\sqrt{\text{MSE}}$ | 95% CI |
| --- | --- | --- | --- | --- | --- | --- | --- | --- | --- | --- | --- | --- |
| Scenario I |  |  |  |  |  |  |  |  |  |  |  |  |
| LM-Naive |  | 1.000 | 0.035 | 0.034 | 0.035 | 94.8 |  | 2.001 | 0.035 | 0.034 | 0.035 | 93.8 |
| PS-Naive |  | 1.000 | 0.035 | 0.041 | 0.035 | 98.0 |  | 2.001 | 0.035 | 0.041 | 0.035 | 98.0 |
| DML-DR |  | 1.029 | 0.036 | 0.037 | 0.046 | 89.3 |  | 2.029 | 0.038 | 0.044 | 0.048 | 94.0 |
| DML-PLM | 1 | 1.001 | 0.034 | 0.035 | 0.034 | 96.3 | 2 | 1.986 | 0.036 | 0.035 | 0.038 | 92.7 |
| semiDNN |  | 0.986 | 0.035 | 0.034 | 0.038 | 92.8 |  | 1.969 | 0.036 | 0.035 | 0.047 | 84.8 |
| deepTL |  | 1.001 | 0.035 | 0.034 | 0.035 | 95.1 |  | 2.002 | 0.035 | 0.034 | 0.035 | 94.2 |
| LM-Oracle |  | 1.000 | 0.035 | 0.034 | 0.035 | 94.8 |  | 2.001 | 0.035 | 0.034 | 0.035 | 93.8 |
| Scenario II |  |  |  |  |  |  |  |  |  |  |  |  |
| LM-Naive |  | 0.587 | 0.059 | 0.059 | 0.417 | 0.0 |  | 1.584 | 0.058 | 0.059 | 0.420 | 0.0 |
| PS-Naive |  | 0.585 | 0.059 | 0.062 | 0.419 | 0.0 |  | 1.582 | 0.058 | 0.062 | 0.422 | 0.0 |
| DML-DR |  | 0.958 | 0.037 | 0.038 | 0.056 | 81.0 |  | 1.958 | 0.040 | 0.045 | 0.058 | 86.7 |
| DML-PLM | 1 | 0.944 | 0.035 | 0.040 | 0.066 | 72.2 | 2 | 1.905 | 0.039 | 0.040 | 0.103 | 33.5 |
| semiDNN |  | 0.987 | 0.036 | 0.035 | 0.038 | 92.3 |  | 1.972 | 0.035 | 0.035 | 0.045 | 88.3 |
| deepTL |  | 1.001 | 0.036 | 0.034 | 0.036 | 93.8 |  | 2.000 | 0.035 | 0.034 | 0.035 | 95.1 |
| LM-Oracle |  | 1.000 | 0.034 | 0.033 | 0.034 | 94.4 |  | 1.999 | 0.032 | 0.033 | 0.032 | 95.2 |
| Scenario III |  |  |  |  |  |  |  |  |  |  |  |  |
| LM-Naive |  | 1.529 | 0.058 | 0.057 | 0.533 | 0.0 |  | 2.528 | 0.059 | 0.057 | 0.532 | 0.0 |
| PS-Naive |  | 1.531 | 0.059 | 0.060 | 0.535 | 0.0 |  | 2.530 | 0.059 | 0.061 | 0.534 | 0.0 |
| DML-DR |  | 1.229 | 0.046 | 0.043 | 0.234 | 0.2 |  | 2.230 | 0.047 | 0.051 | 0.235 | 0.2 |
| DML-PLM | 1 | 1.141 | 0.040 | 0.044 | 0.146 | 7.5 | 2 | 2.109 | 0.041 | 0.044 | 0.116 | 29.2 |
| semiDNN |  | 0.987 | 0.036 | 0.036 | 0.039 | 92.7 |  | 1.971 | 0.038 | 0.037 | 0.048 | 86.7 |
| deepTL |  | 0.998 | 0.036 | 0.036 | 0.036 | 94.6 |  | 1.997 | 0.038 | 0.036 | 0.038 | 93.8 |
| LM-Oracle |  | 0.999 | 0.031 | 0.032 | 0.031 | 94.9 |  | 1.999 | 0.032 | 0.032 | 0.032 | 94.9 |

In Scenario II, as complex structures are introduced in *e*(***X***) and *γ*(***X***), naive methods, including LM-Naive and PS-Naive, fail as expected due to violations of the model assumptions. DML-PLM and DML-DR both show a reduced bias and Monte Carlo standard error, compared to naive methods. However, the biases from these methods are still not minor, leading to poor 95% CI coverages. semiDNN outperforms these methods. As expected, deepTL outperforms semiDNN especially when *β* = 2, with an ignorable bias and a smaller standard error that is only slightly larger than that of LM-Oracle.

In Scenario III, deepTL and semiDNN continue to achieve a better performance than other competing methods, further indicating the robustness and advantages of our proposed methods. deepTL continues to perform similarly as LM-Oracle, with a significant improvement in bias compared to semiDNN. As more complex structure is introduced, DML-PLM and DML-DR show more severe biases and worse 95% CI coverages, and LM-Naive and PS-Naive continue to show a remarkably large bias.

To further investigate the performance of the proposed methods, we extend the simulations with different *N, p, σ* and *β* values under the same setting of Scenario III. Each time we vary one of the parameters and keep the rest of the parameters fixed. We exclude the results for LM-Naive and PS-Naive due to their poor performance. The results are displayed in Figure 1.

**Fig 1.**
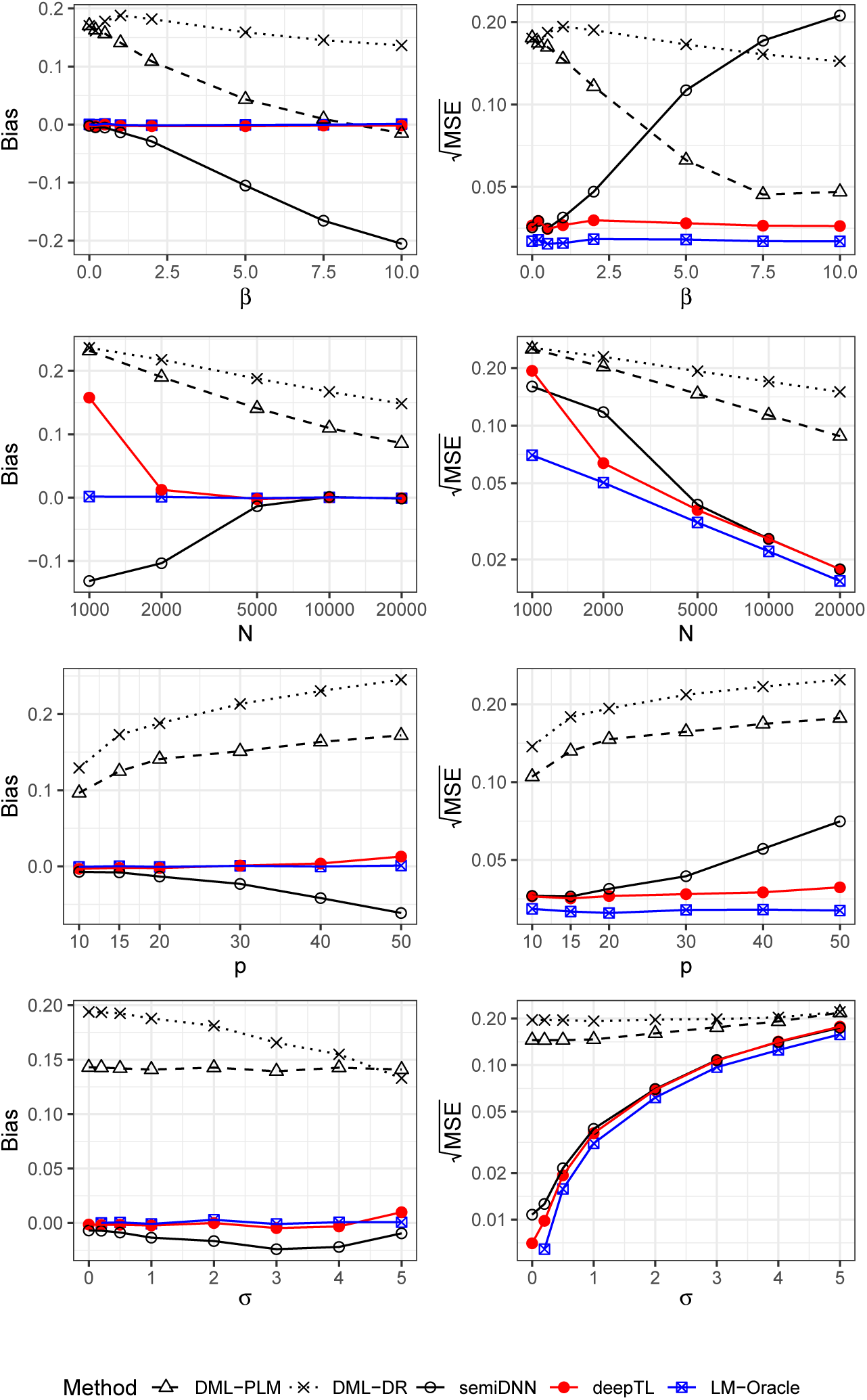
The bias and MSE plot for the simulation setting in Scenario III.

First, we observe that the bias from semiDNN increases with the increment of *β*. In contrast, deepTL consistently offers an unbiased treatment effect estimate, regardless of the true treatment effect value. Next, as we increase *N* from 1,000 to 20,000, as expected, all methods result in reduced biases and MSE. deepTL shows a negligible bias as *N* increases to 2,000 or above, while the other methods still show noticeable biases at *N* = 2000. Moreover, as *p* increases from 10 to 50, semiDNN, DML-PLM and DML-DR all show increased biases, while deepTL is not affected as severely as the others. Finally, deepTL and semiDNN outperform DML-PLM and DML-DR regardless of the value of *σ*. In summary, deepTL performs better than, or equally as well as other methods across all simulation parameters, reflecting its advantages with the modification of the semiparametric framework procedure and the use of bagged DNN models.

Additionally, to investigate how the model size, including *L* and *n*_*l*_ (*l* = 1, …, *L*), and the number of DNNs, *K*, affect the performance of deepTL, more simulations under the setting of Scenario III for *β* = 1 are conducted. The results are presented in Figure 2. In the top plots, bagged DNNs have fixed *K* = 100 but *L* ranges from 1 to 6, *n*_1_ ranges from 10 to 100 as the x-axis, 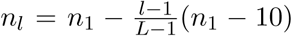 for *l* = 2, …, *L*, while in the bottom plots, bagged DNNs are fixed with *L* = 6, (*n*_1_, …, *n*_6_) = (20, 18, 16, 14, 12, 10) but *K* varies from 2 to 500.

**Fig 2.**
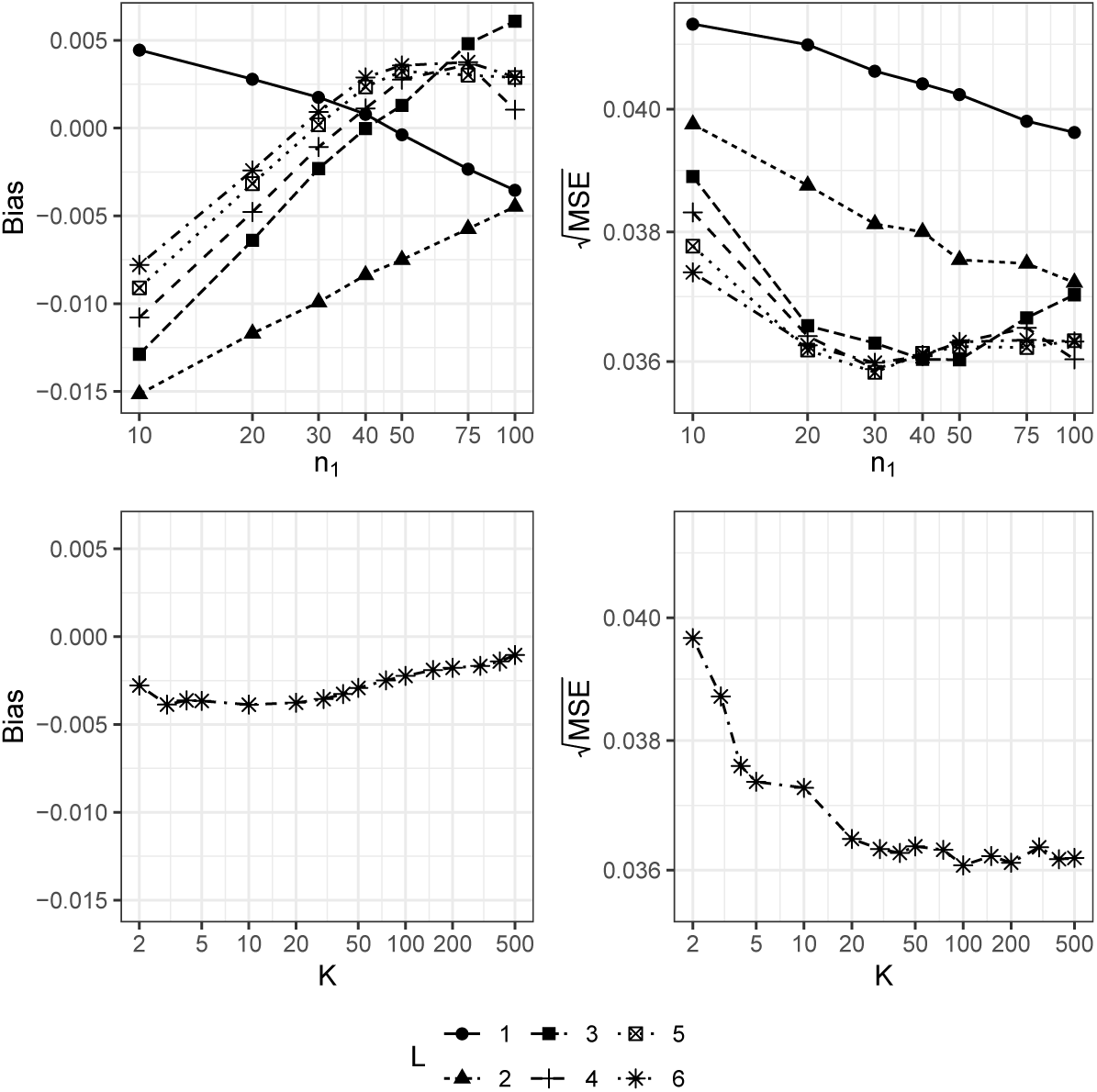
The bias and MSE plot for deepTL using different L, n1 and K in bagged DNNs for the simulation setting in Scenario III when β = 1.

Clearly, the impacts of *L, n*_1_ and *K* on deepTL are minor. In the worst case where *L* = 2, *n*_1_ = 10 and *K* = 100, the absolute bias is 0.015, smaller than those from other competing methods, while for most cases, the absolute bias of deepTL is less than 0.005. For MSE, we have similar observations. deepTL with *n*_1_ ∈ [20, 50] and *L* ≥ 3 in general performs well. Another encouraging finding is that the performance of deepTLincreases with the increase of *K*. However the improvement becomes less obvious when *K* > 100, suggesting that deepTL only requires a moderate number of bagged DNNs.

### 3.2. A Post-surgery Pain Study

To illustrate the practical application of deepTL, we apply it to a post-surgery pain data set obtained from the University of Florida Integrated Data Repository, a large-scale EHR database.

The study (Tighe et al., 2016) included 3196 patients who underwent different surgeries related to the digestive system, the musculoskeletal system and the nervous system. One of the objectives of the study was to compare two anesthetic procedures, i.e. nerve block (*Z* = 1) versus general anesthesia (*Z* = 0), on relieving the severest post-surgery pain intensity within 72 hours after the surgery. Among these patients, 2438 (76.3%) patients chose the nerve block procedure while the remaining patients opted for general anesthesia. The nerve block procedure interrupts signals traveling along a nerve and is often used for pain relief. Compared to the traditional anesthesia procedure, the nerve block has some advantages by allowing patients to remain awake, thereby avoiding some adverse reactions of general anesthesia, such as cognitive loss. However, it is clinically important to test if the nerve block procedure is as effective as the general anesthesia. The primary outcome of the study was that the severest post-surgery pain intensity was experienced by patients within 72 hours after the surgery, which is quantified numerically and scaled between 0 and 10, where the higher pain scores mean more severe pain.

Covariates other than the treatment procedures (i.e. nerve block and general anesthesia) include patient age, gender, ethnicity, body mass index (BMI), surgical duration, marital status, opioid use, muscle relaxant use, nonsteroidal anti-inflammatory drugs (NSAIDs) use, benzo use, selective serotonin reuptake inhibitors (SSRIs) use, and current procedural terminology (CPT). Distributions of the baseline covariates by treatment group are presented in Table 2. We first apply an ANOVA method to the postoperative pain data and compare the effects of the two pain relief methods without any adjustment of the covariates. We also employ methods described in Section 3.1. The results are summarized in Table 3.

**Table 2.** Baseline Distribution of the Post-surgery Study Data.

|  |  | Nerve Block | General |
| --- | --- | --- | --- |
|  |  | 2438 (76.3%) | 758 (23.7%) |
|  |  | Mean (SD) | Mean (SD) |
| Maximum pain intensity |  | 8.0 (2.6) | 8.2 (2.4) |
| Age (year) |  | 58.1 (15.1) | 55.4 (16.1) |
| BMI (kg/m<sup>2</sup>) |  | 29.6 (7.7) | 29.2 (8.3) |
| Surgical Duration (hour) |  | 3.9 (2.1) | 3.1 (2.0) |
|  |  | N (%) | N (%) |
| Gender | Male | 1158 (47.5%) | 360 (47.5%) |
|  | Female | 1280 (52.5%) | 398 (52.5%) |
| Ethnicity | Non-Hispanic | 2277 (93.4%) | 716 (94.5%) |
|  | Hispanic | 49 (2.0%) | 26 (3.4%) |
|  | Unknown | 112 (4.6%) | 16 (2.1%) |
| Marital | Married | 1329 (54.5%) | 381 (50.3%) |
|  | Single | 602 (24.7%) | 225 (29.7%) |
|  | Other | 507 (20.8%) | 152 (20.1%) |
| Opioid | Yes | 977 (40.1%) | 296 (39.1%) |
|  | No | 1461 (59.9%) | 462 (60.9%) |
| Muscle relaxant | Yes | 204 (8.4%) | 49 (6.5%) |
|  | No | 2234 (91.6%) | 709 (93.5%) |
| NSAID | Yes | 1418 (58.2%) | 409 (54.0%) |
|  | No | 1020 (41.8%) | 349 (46.0%) |
| Benzo | Yes | 272 (11.2%) | 89 (11.7%) |
|  | No | 2166 (88.8%) | 669 (88.3%) |
| SSRI | Yes | 609 (25.0%) | 182 (24.0%) |
|  | No | 1829 (75.0%) | 576 (76.0%) |
| CPT count | 0 – 2 | 1952 (80.1%) | 620 (81.8%) |
|  | 3 – 4 | 433 (17.8%) | 127 (16.8%) |
|  | ≥ 5 | 53 (2.2%) | 11 (1.5%) |

**Table 3.** Post-operative Pain Analysis.

| Method | $\hat{\beta}$ | $\widehat{SE}(\hat{\beta})$ | p-value |
| --- | --- | --- | --- |
| ANOVA | -0.180 | 0.105 | 0.086 |
| LM-Naive | -0.200 | 0.106 | 0.060 |
| PS-Naive | -0.205 | 0.107 | 0.056 |
| DML-DR | -0.079 | 0.218 | 0.718 |
| DML-PLM | -0.138 | 0.105 | 0.188 |
| semiDNN | -0.221 | 0.108 | **0.040** |
| deepTL | -0.222 | 0.108 | **0.040** |

The first row of Table 3 presents the crude treatment effect estimate, i.e. −0.18, obtained from ANOVA. The nerve block is not significantly different from general anesthesia in this analysis (*p* = 0.086). A similar conclusion is obtained from LM-Naive and PS-Naive, even after the confounding covariates are adjusted, i.e. there exists no significant difference between the two comparison procedures. Besides, DML-DR and DML-PLM show a less significant result. In contrast, the result from deepTL demonstrates that at the 0.05 significant level, the nerve block procedure is significantly more effective in relieving the severest post-surgery pain intensity than the general anesthesia procedure does (*p* = 0.04). This conclusion is also consistent with the results of earlier clinical studies (Tverskoy et al., 1990; Shir, Raja and Frank, 1994).

## 4. Discussion

In this paper, we proposed a powerful DNN-based semiparametric framework, deepTL, to adjust the complex confounding structures in comparative effectiveness analysis. As a universally consistent approximator, DNN has a unique advantage over parametric supervised learning methods, as well as other non-parametric machine learning approaches. In addition, bagging and the proposed ensembling scheme reduce the level of overfitting and increase the accuracy of DNN approximating functions, which consequently increases the treatment effect estimate accuracy (see Table S1 of Supplement B). Though the estimator from semiDNN enjoys root-N-consistency, we have shown that the method can have an elevated bias under finite sample settings when the underlying treatment effect *β* is not small which motivates the development of deepTL. Extensive simulation studies demonstrate that deepTL consistently outperforms other existing competing methods for data with complex confounding structures, while under simple settings, deepTL could still perform as well as other competing methods.

For observational data, in addition to confounding bias, overadjustment bias is another concern (Schisterman, Cole and Platt, 2009). To study overadjustment bias of deepTL, we carefully designed a new simulation setup, similar to DAG2 in Schisterman, Cole and Platt (2009). Under DAG2, we found that overadjustment bias is common to all comparing methods, while deepTL delivers the closest estimate to that from LM-Oracle (see Table S2 in Supplement B). Furthermore, when there exist unmeasured confounding factors, the treatment effect estimate from deepTL, as well as other existing methods, can be biased in general. However, deepTL appears to control the bias to the minimal (see Table S3 of Supplement B for simulation results on studying the robustness of deepTL on data with unmeasured confounding factors).

In deepTL, for computational efficiency, we set *β*_1_ to the estimate from the PS covariate adjustment method which works sufficiently well in improving the performance of deepTL over semiDNN. Alternatively, one could set *β*_1_ to the estimate from semiDNN which is however, more computationally demanding.

The proposed method is developed for continuous outcomes. However, it is not necessary for the DNN to be limited to this data type, though its performance for other data types, e.g. binary outcomes, deserves further investigation. Furthermore, EHR data is often clustered and with repeated measurements from the same individual. How to extend the proposed method to non-independent observational data is practically important as well, but is beyond the scope of this paper.

## Acknowledgement

We thank the Editor and anonymous reviewers for their insightful comments and constructive suggestions, which have greatly improved the paper.

## SUPPLEMENTARY MATERIAL

### Supplement A: R package “deepTL”

(https://github.com/SkadiEye/deepTL). We have developed an R package, Deep Treatment Learning “deepTL”, to implement our proposed method.

### Supplement B: Additional simulation studies

(doi: COMPLETED BY THE TYPESETTER; .pdf). We provide additional supporting simulation studies to illustrate (i) the scenarios with unmeasured confounding; (ii) necessity of the bagging procedure; (iii) *β*_1_ estimated by semiDNN.

