## Supplement for "A Deep Learning Semiparametric Regression for Adjusting Complex Confounding Structures"

### More results with Scenario III

In this section, more simulations in Scenario III are conducted to investigate how the performance of deepTL and/or semiDNN is affected by: i) using a single DNN, denoted as deepTL (S-DNN) and semiTL (S-DNN), respectively, corresponding to deepTL and semiDNN in the paper; ii) using bagged DNNs without the proposed score-based filtering procedure, denoted as deepTL (B-DNN) and semiDNN (B-DNN).

As shown in Table S1 below, it clearly indicates that bagging is helpful as the parameter estimate has a notably larger RMSE from the models without bagging than those from the models with bagging. The reason to develop the bootstrap aggregation scheme (i.e. bagging with score-based filtering procedure) is motivated by the discussion in Zhou et al. (2002) as well as our own observation (Mi et al., 2019) that under certain circumstances, in particular when the number of covariates is large, some DNNs in the ensemble model may not perform as well as others, and our bootstrap aggregation scheme could potentially filter "bad" DNNs out and thus potentially benefits the final ensemble, which is demonstrated by the results in Table S1 (i.e. deepTL).

| Method | $\beta$ | Avg($\widehat{\beta}$) | SE($\widehat{\beta}$) | $\widehat{\text{SE}}$($\widehat{\beta}$) | $\sqrt{\text{RMSE}}$ |
|---|---|---|---|---|---|
| semiDNN (S-DNN) | 1 | 0.980 | 0.046 | 0.040 | 0.050 |
| semiDNN (B-DNN) | | 0.989 | 0.037 | 0.038 | 0.039 |
| deepTL (S-DNN) | | 0.993 | 0.045 | 0.040 | 0.045 |
| deepTL (B-DNN) | | 0.992 | 0.037 | 0.038 | 0.038 |
| deepTL | | 0.998 | 0.036 | 0.036 | 0.036 |
| semiDNN (S-DNN) | 2 | 1.966 | 0.067 | 0.040 | 0.075 |
| semiDNN (B-DNN) | | 1.983 | 0.038 | 0.039 | 0.041 |
| deepTL (S-DNN) | | 1.992 | 0.045 | 0.040 | 0.045 |
| deepTL (B-DNN) | | 1.992 | 0.037 | 0.038 | 0.038 |
| deepTL | | 1.997 | 0.038 | 0.036 | 0.038 |

Table S1: Scenario III with additional methods.

### Overadjustment Bias

In causal inference, Schisterman et al. (2009) have raised an awareness of overadjustment bias, caused by control of (unnecessary) intermediate variables. In this section, additional simulations on this specific issue are performed to illustrate the performance of deepTL under such scenarios. Specifically, we simulate the situation described by the causal diagram in Figure S1 (simliar to DAG2 of Schisterman et al. (2009)) where

$$\log \frac{e(\boldsymbol{X})}{1-e(\boldsymbol{X})} = 1 - \sum_{j=1}^{5} \alpha_{1,j} X_j - X_8^2$$
$$Y = \alpha_Z Z + \alpha_Y U - 1 + \sum_{j=1}^{5} \alpha_{2,j} X_j + |X_8| X_9^2 + \epsilon_Y.$$

Here $U$ is an unmeasured intermediate variable that is dependent on $Z$, or $U = \alpha_U(Z - 0.5) + \epsilon_U$, $M$ is an observed variable that depends on $U$, or $M = \alpha_M U + \epsilon_M$, $\boldsymbol{X} \sim \text{MVN}(\mathbf{0}, I_p)$ and $\epsilon_Y, \epsilon_U, \epsilon_M \sim \text{N}(0, 1)$. We set $N$, $p$, $\boldsymbol{\alpha}_1$ and $\boldsymbol{\alpha}_2$ to be the same as those in Scenario III, $\alpha_Z = 1$ or 2 in different scenarios, $\alpha_Y = 1$, $\alpha_U = 0.25$ and $\alpha_M = 0.5$, which results in the total treatment effect $\beta = \alpha_Z + \alpha_Y \alpha_U = \alpha_Z + 0.25$. A model that adjusts for $M$ can lead to the so called overadjustment bias

$$\text{Bias}_\text{M} = \frac{\alpha_Y \alpha_U}{1 + \alpha_M^2} - \alpha_Y \alpha_U$$

defined in (Schisterman et al., 2009), which equals -0.05 for the simulation.

The simulation results are displayed in Table S2. For all the methods compared, $U$ is assumed to be unobserved, including LM-Oracle. If only $X$ but not $M$ are used for adjusting the confounding effects, LM-Oracle has a negligible bias with the smallest RMSE, followed by deepTL, while other methods are more significantly biased. However, if $M$ is included as an additional covariate, all methods now become biased. The bias of LM-Oracle is about $-0.05$ which is close to the expected overadjustment bias. Again, deepTL provides the closest estimate of $\beta$ to that of LM-Oracle.

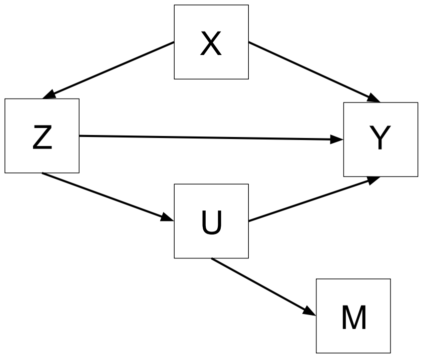


Figure S1: Causal diagram for simulation scenario with overadjustment bias

| Method | $\beta$ | Avg($\widehat{\beta}$) | SE($\widehat{\beta}$) | $\widehat{\text{SE}}(\widehat{\beta})$ | $\sqrt{\text{RMSE}}$ |
|---|---|---|---|---|---|
| | | Exclude $M$ | | | |
| LM-Naive | | 1.778 | 0.067 | 0.065 | 0.532 |
| PS-NAive | | 1.780 | 0.067 | 0.068 | 0.534 |
| DML-DR | | 1.479 | 0.056 | 0.053 | 0.236 |
| DML-PLM | 1.25 | 1.381 | 0.051 | 0.055 | 0.140 |
| semiDNN | | 1.231 | 0.050 | 0.051 | 0.053 |
| deepTL | | 1.242 | 0.050 | 0.051 | 0.050 |
| LM-Oracle | | 1.248 | 0.044 | 0.045 | 0.044 |
| LM-Naive | | 2.778 | 0.067 | 0.065 | 0.532 |
| PS-NAive | | 2.780 | 0.067 | 0.068 | 0.534 |
| DML-DR | | 2.479 | 0.056 | 0.060 | 0.235 |
| DML-PLM | 2.25 | 2.349 | 0.051 | 0.055 | 0.111 |
| semiDNN | | 2.218 | 0.050 | 0.051 | 0.059 |
| deepTL | | 2.242 | 0.050 | 0.051 | 0.050 |
| LM-Oracle | | 2.248 | 0.044 | 0.045 | 0.044 |
| | | Include $M$ | | | |
| LM-Naive | | 1.728 | 0.066 | 0.063 | 0.482 |
| PS-NAive | | 1.730 | 0.066 | 0.068 | 0.484 |
| DML-DR | | 1.455 | 0.054 | 0.052 | 0.212 |
| DML-PLM | 1.25 | 1.350 | 0.049 | 0.053 | 0.112 |
| semiDNN | | 1.179 | 0.047 | 0.049 | 0.085 |
| deepTL | | 1.191 | 0.047 | 0.049 | 0.076 |
| LM-Oracle | | 1.198 | 0.042 | 0.042 | 0.067 |
| LM-Naive | | 2.728 | 0.066 | 0.063 | 0.482 |
| PS-NAive | | 2.730 | 0.066 | 0.068 | 0.484 |
| DML-DR | | 2.455 | 0.054 | 0.059 | 0.212 |
| DML-PLM | 2.25 | 2.312 | 0.050 | 0.053 | 0.079 |
| semiDNN | | 2.165 | 0.048 | 0.049 | 0.097 |
| deepTL | | 2.191 | 0.047 | 0.049 | 0.076 |
| LM-Oracle | | 2.198 | 0.042 | 0.042 | 0.067 |

Table S2: Simulation results for overadjustment bias

### Unmeasured Confounding

Similar to the PS approaches (Rosenbaum and Rubin, 1983), one assumption underlying the proposed method is that the confounding factors are fully observed. However, in practice, some confounding factors might be unobserved. To investigate how deepTL behaves with unmeasured confounding factors, we conduct further simulation studies where the true model is similar to Scenario III,

$$\log \frac{e(\boldsymbol{X})}{1 - e(\boldsymbol{X})} = 1 - \sum_{j=1}^{5} \alpha_{1,j} X_j - X_8^2,$$

$$\gamma(\boldsymbol{X}) = -1 + \sum_{j=1}^{5} \alpha_{2,j} X_j + |X_8| X_9^2,$$

where $\boldsymbol{X} \sim \text{MVN}(\mathbf{0}, \Sigma)$, $\Sigma = \{\sigma_{jj'}\}_{1 \le j,j' \le p}$, $\sigma_{jj'} = 1$ for $j = j'$ and 0.5 otherwise with $X_1, ..., X_5$ being unmeasured confounding factors, that is, they are assumed to be unavailable for all but the oracle method. We set $N$, $p$, $\beta$, $\boldsymbol{\alpha}_1$ and $\boldsymbol{\alpha}_2$ to be the same as those in Scenario III.

The results are presented in Table S3. Due to the presence of unmeasured confounding, the bias increases inevitably, for all methods except LM-Oracle. In this scenario, both semiDNN and deepTL are less biased, compared to the other methods. Moreover, deepTL continues to outperform semiDNN.

| Method | $\beta$ | Avg($\widehat{\beta}$) | SE($\widehat{\beta}$) | $\widehat{\text{SE}}$($\widehat{\beta}$) | $\sqrt{\text{RMSE}}$ |
|---|---|---|---|---|---|
| LM-Naive | | 1.901 | 0.080 | 0.073 | 0.905 |
| PS-Naive | | 1.885 | 0.078 | 0.074 | 0.888 |
| DML-DR | | 1.247 | 0.060 | 0.056 | 0.254 |
| DML-PLM | 1 | 1.116 | 0.039 | 0.041 | 0.123 |
| semiDNN | | 1.076 | 0.037 | 0.040 | 0.085 |
| deepTL | | 1.071 | 0.037 | 0.040 | 0.080 |
| LM-Oracle | | 0.999 | 0.032 | 0.032 | 0.032 |
| LM-Naive | | 2.901 | 0.080 | 0.073 | 0.905 |
| PS-Naive | | 2.886 | 0.079 | 0.074 | 0.890 |
| DML-DR | | 2.247 | 0.059 | 0.061 | 0.254 |
| DML-PLM | 2 | 2.105 | 0.040 | 0.041 | 0.112 |
| semiDNN | | 2.078 | 0.038 | 0.041 | 0.086 |
| deepTL | | 2.071 | 0.037 | 0.040 | 0.080 |
| LM-Oracle | | 1.999 | 0.032 | 0.032 | 0.032 |

Table S3: Simulation results for unmeasured confounding.
